# Uncertainty revealed by delayed responses during olfactory matching

**DOI:** 10.1101/2022.09.11.507462

**Authors:** Rajdeep Bhowmik, Meenakshi Pardasani, Sarang Mahajan, Anindya S. Bhattacharjee, Sasank Konakamchi, Shambhavi Phadnis, Thasneem Musthafa, Eleanor McGowan, Priyadharshini Srikanth, Shruti D. Marathe, Nixon M. Abraham

## Abstract

Matching of olfactory stimuli involves both sensory and higher cognitive functioning. Different decision processes such as detection and discrimination, along with holding the perceived information are involved during the matching process. Accuracy and decision times, the interdependent readouts, can define the uncertainty involved in matching of sensory stimuli. To probe sensory and cognitive functions involving olfactory system in human subjects, we have developed a novel olfactory matching paradigm using an automated custom-built olfactory-action meter. With precise and consistent odor delivery and real-time data analysis, our system automates the entire process without any intervention by the experimenter, making it suitable as a diagnostic tool for quantifying olfactory and neurocognitive fitness. In around 400 healthy human subjects, with mean detection accuracy of 90%, we observed significantly better olfactory matching performance for simple monomolecular odors, in comparison to complex binary odor mixtures. Odor matching accuracy declined significantly with the increase in odor complexity. Olfactory matching was more rapid when subjects made correct versus incorrect decisions, indicating perceptual certainty. Subjects also took longer matching time for complex odors compared to simple odor stimuli. Thus, olfactory matching that provides a combined readout of sensory and cognitive fitness, establishes a direct link between the performance accuracy and the certainty of decisions.

## INTRODUCTION

A fundamental challenge for the brain is encoding specific sensory information for variable durations to facilitate discriminations in a complex sensory environment. Decisions with certainty in response to perceived information would involve successful detection of the stimuli followed by weighing sensory evidences, which aid in generating a motor action to execute the formed decision [1,2]. Psychophysical readouts of accuracy and time needed for decisions, which are inter-dependent parameters, equip us to quantitatively measure this certainty [3–6]. Visual dot motion discriminations and auditory time-varying directional cues are often utilized to detail how subjects’ performance accuracy and decision times can reveal the concepts underlying uncertainty of decision-making and concurrently explore the effect of stimulus-complexity [7–9]. However, it remains elusive how the variable time delays between subsequent sensory stimuli affect the accuracy, and certainty of decisions in an odor-based decision-making task. To probe this, we decided to carry out olfactory matching in a variable time-elapsed task using a custom-built olfactory-action meter (OAM) across different stimulus complexities.

Over the past few decades, many psychophysical and psychosocial studies have investigated olfactory capabilities in humans [10]. Interest to try different behavioral tests and readouts assessing olfactory capabilities in humans grew as a result of the possible correlation of deficits in sense of smell with certain brain disorders. Olfactory impairments are observed in patients with neurodegenerative disorders at the early onset of diseases [11,12,13]. Olfactory functions are also negatively affected in patients with certain traumatic brain injuries [14]. More prominently, cases of anosmia (loss of sense of smell) and hyposmia/microsmia (weakened sense of smell) have been reported in many patients suffering from CoronaVirus disease-2019 (COVID-19) [15–19]. Therefore, quantifying olfactory fitness is extremely valuable in diagnosing many neurodegenerative and infectious diseases.

Most psychophysical studies in the field utilize semantic and/or non-semantic, analytical tests. These include Sniffing stick, University of Pennsylvania Smell Identification test (UPSIT), Scandinavian odor identification test, two-alternative forced choice odor detection threshold test etc [13,20,21]. However, olfactory dysfunctions of low severity can go unnoticed due to excessive dependence on subjective evaluations [22]. Therefore, novel approaches combining sensitive measurements of olfactory thresholds with objective evaluations would result in better diagnosis of olfactory dysfunctions [16,23]. Further, to use olfaction for quantifying cognitive impairments, introduction of working memory functions would be expedient. Hence, we have carried out olfactory matching tests where each participant needs to decide whether the presented odors with a specified delay were same or different. The complexity of stimuli was increased by delivering binary odor mixtures. While we observed an average detection accuracy of 90% for both simple and complex odors, better olfactory matching accuracy was observed for monomolecular odor pairs compared to binary mixtures. To correlate the certainty of their decisions with accuracy, we calculated olfactory matching times for correct and incorrect responses. We find that correct responses were registered faster compared to incorrect ones. Also, subjects were slower in correctly matching the binary mixtures as compared to monomolecular odors. By incorporating different decision processes, detection and discrimination along with working memory functions, our method offers an ideal tool for quantifying sensory and cognitive deficits.

## RESULTS AND DISCUSSION

Psychophysical tasks and tools equip us to quantitatively measure human perception across different sensory and cognitive domains [6]. Questionnaire based methods allow for sensory stimulus/object detection assessment in humans [24]. To begin with, we instructed the participants to smell odorous objects available in their households and fill their detection responses in an online questionnaire (see Methods). We assigned a 1 to the detected odorous object while 0 was assigned if they could not smell it. Detection indices were calculated based on the average of 143 subject responses across 33 odorous objects belonging to different clusters (Figure 1A). This experimental approach had it’s short-comings: no control over odor delivery without any absolute time onset and offset of emitted smell; the odorous objects consist of complex mix of several monomolecular odorants; and some of the objects were also prone to over-ripening or losing their characteristic smell due to variations of environmental factors in different households across the country. Such a method, thus cannot facilitate revealing olfactory perceptual changes in humans with high precision. In order to reliably assess the sensory-cognitive fitness using olfaction in humans, we developed an Olfactory-action meter (OAM) [17] (Figure 1B). In this system, odor (either monomolecular or binary mixture synthetic odor chemicals diluted in mineral oil) delivery is automated with a precise control over its time onset and offset as revealed by the photo-ionization detection (PID) measurements (Figure 1C). Irrespective of the intrinsic differences in the physico-chemical properties of odors, our instrument delivered the odor with a sharp pulse for a stimulus presentation time of 4s. The odor-controlling unit contained 10 mass flow controllers, which independently regulated air delivery through odor reservoirs. The output from these reservoirs was further diluted with a stream of clean air. The odor-air mixture was then delivered through a 12-channel glass funnel (Figure 1B). The output from the funnel was attached to a ventilator mask. At the beginning of each session, subjects were evaluated for their odor detection abilities. We found out high and similar detection indices (> 0.8) for 9 monomolecular odorants delivered through OAM across 180 subjects (Figure 1D).

**Figure 1:**
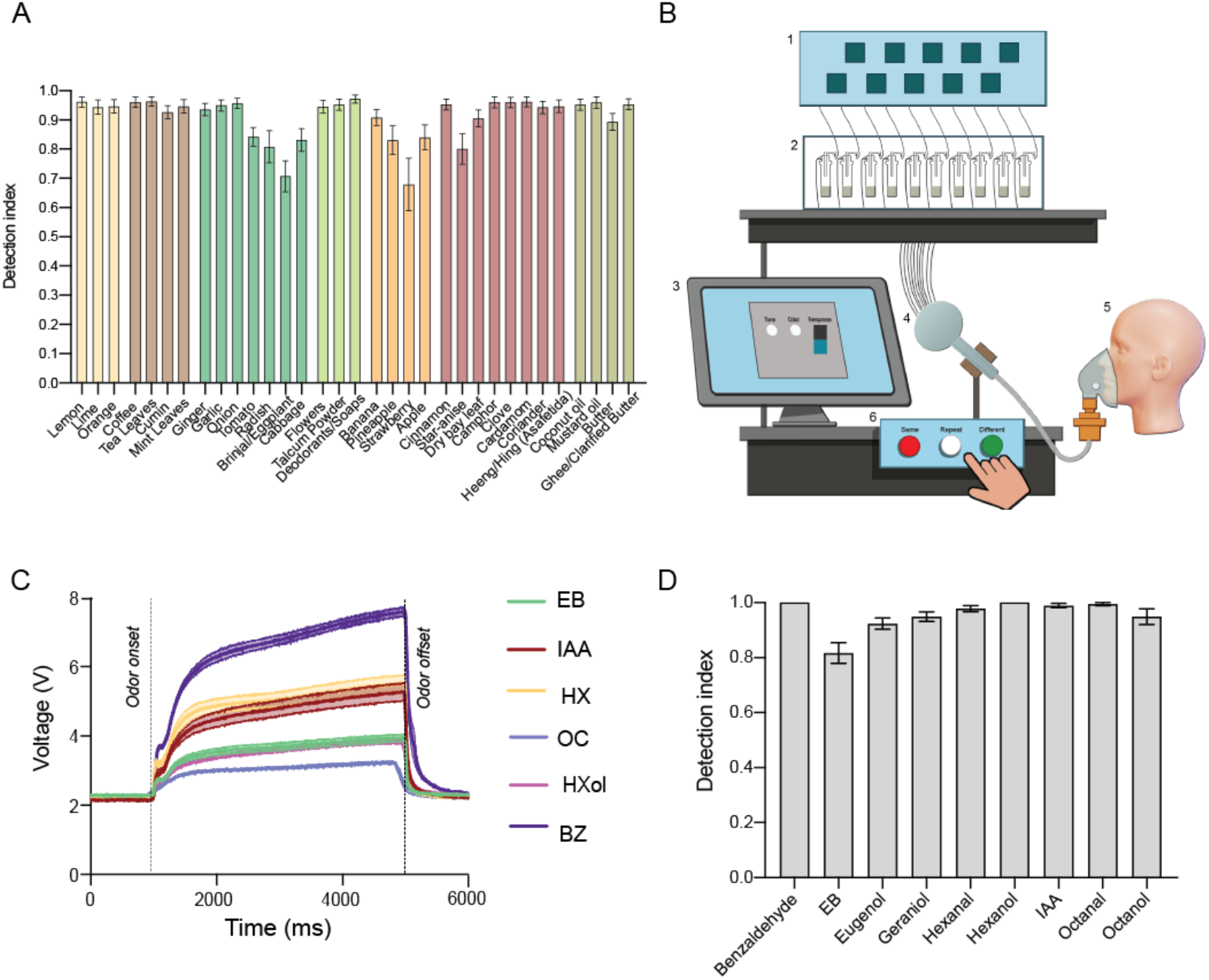
Development of an Olfactory-action meter for quantifying olfactory fitness in humans. A. Average detection indices (DI) for commonly occurring household odorous objects and food items in humans. Data was collected using an online questionnaire to probe the smell detection abilities of humans. For each subject, DI value of ‘1’ indicated successful detection while ‘0’ meant failed at detecting the object using olfaction. Odorous items were divided into Clusters; Cluster 1 composed of ‘Citrus fruits’ (Yellow bars), Cluster 2 had ‘Odorous food items’ (Brown and Green bars), Cluster 3 encompassing flowers, talcum and deodorants/soaps (Light green bars), Cluster 4 consisted of ‘fruits other than citrus’ (Orange bars), Cluster 5 included ‘spices’ (Maroon bars) and Cluster 6 consisted of ‘Fats and Oils’ (Olive colored bars) (N = 143 subjects). B. Diagram depicting the custom-built olfactory action-meter (OAM). 1. Flow manifold, 2. Odorous bottles consisting of diluted synthetic odors, 3. Computer monitor, 4. Glass funnel-shaped nozzle, 5. Human with a ventilating mask for odor delivery, 6. Response console to register the response in an Olfactory Matching (OM) task. Upon initiation of a trial in an OM task, flowmeters delivered air to the odor bottles. Odorized air was mixed with fresh air and passed through a 12-channel glass funnel. Odor output from the funnel, via a nozzle, delivered odorized air to the ventilator mask worn by the human subject via a tube. After the onset of the second odor, the participant could respond by pressing on one of the three buttons of the response console, i.e, *Same* (the two odors were perceived as same), *Different* (the two odors were perceived as different) or *Repeat* (judgement could not be made and the trial is repeated), kept in front of the human performing the task. C. Photoionization detection (PID) measurements of changes in voltage amplitudes for simple odors are plotted. Odor was delivered for 4s. Values were averaged across 10 sets of measurements per odor. D. Detection indices of odors delivered via the OAM, averaged across subjects (N = 180 subjects, average value of DI varies from 0.81 to 1 suggesting that > 80% of the subjects were able to detect all the odors).

The odors detected by the subjects during the detection task were then selected for Olfactory Matching (OM) task. For a particular trial of an OM task, the subject had to sample and compare two odors delivered and provide a response of whether the two odor stimuli were ‘same’ or ‘different’ (See trial workflow in Supplementary figure 1) by pressing on the response console placed in front of the subject (Figure 1B). It is a convenient, easy to understand console with three color-coded response option buttons (same, different or repeat). A display screen placed in front of the subject provided a cue to notify the odor delivery onset and a real time response timer to indicate the response window. Trial onset was marked by a 200ms tone to alert the individual about the commencement of a new trial [16]. During the pre-COVID-19 era, we carried out OM task to assess working memory (WM) of healthy subjects across different delays between the odors or inter-stimulus intervals (ISI), which ranged from 2.5s to 10s (Figure 2A). By designing our OM task with varying ISIs, we wanted to probe how the neural noise linked to working memory due to long delays affected the uncertainty associated with olfactory matching. A total of 180 subjects (N_2.5s_ = 75 sessions, N_5s_ = 78, N_7.5s_ = 53 and N_10s_ = 63) performed the OM task paradigm. We found that the different ISI durations did not influence the average performance accuracies (Figure 2B; p = 0.42, F = 0.93, One-Way ANOVA). OM task consisted of a random number of both the ‘same’ trials wherein both the presented odors were same and ‘different’ trials wherein the presented odors were different. The differences did not exist either for same or different trials (Figure S2 A1 & A2, One-way ANOVA with Tukey’s Multiple Comparison Test, p > 0.05). We also quantified Olfactory Matching Time (OMT), which is the average time taken by the subject to respond by pressing on the response console in a particular session (averaged across 20 trials). OMT remained comparable across the OM tasks of varying ISI suggesting that the overall response time remains unperturbed by the time lag between the stimuli, at least, with the durations that we tried (Figure 2C; N_2.5s_ = 75 session, N_5s_ = 78, N_7.5s_ = 53 and N_10s_ = 63; p = 0.21, F = 1.524, One-Way ANOVA). This is indeed suggestive of efficient maintenance of olfactory WM across the differing ISIs that we evaluated. Upon separately quantifying accuracy for the two kinds of trials, we found out similar performance levels (Figure 2D; N_2.5s_ = 75 sessions, p = 0.17; N_5s_ = 78, p = 0.36; N_7.5s_ = 53, p = 0.24; and N_10s_ = 63, p = 0.85; Paired t-test). To assess if the perceivable shifts in decision-making across different ISIs for same vs. different stimulus types for correct and incorrect trial responses were occurring, we assessed OMT parameter as well. Across the four ISI conditions, the normalized changes in OMT are found out to be significantly longer for the incorrect trials (Figure 2E, p_2.5s_ < 0.0001, p_5s_ = 0.003, p_7.5s_ < 0.0001 and p_10s_ < 0.0001, Paired t-test for each ISI plot). This confirmed the certainty of decisions with accurately responding trials. Upon considering correct responses, change in OMT between same and different odor pair matching showed shorter time taken to decide whether the matched odors were different as compared to the decision pertaining to same correct decisions, revealing uncertainty involved in matching same stimuli (Figure 2F; p_2.5s_ = 0.0003, p_5s_ < 0.0001, p_7.5s_ = 0.0003 and p_10s_ = 0.0013, Paired t-test for each ISI plot). On the other hand, OMT did not differ between the same and different trial types when only incorrectly responded trials were taken into account (Figure 2G; p_2.5s_ = 0.34, p_5s_ = 0.57, p_7.5s_ = 0.58 and p_10s_ = 0.36, Paired t-test for each ISI plot). This suggests that humans are more accurate when they are responding faster in an OM task, irrespective of any ISI condition. Indeed, OMT is lying between 3s to 5.5s for approximately 80% of the correct responses (Figure 2H).

**Figure 2:**
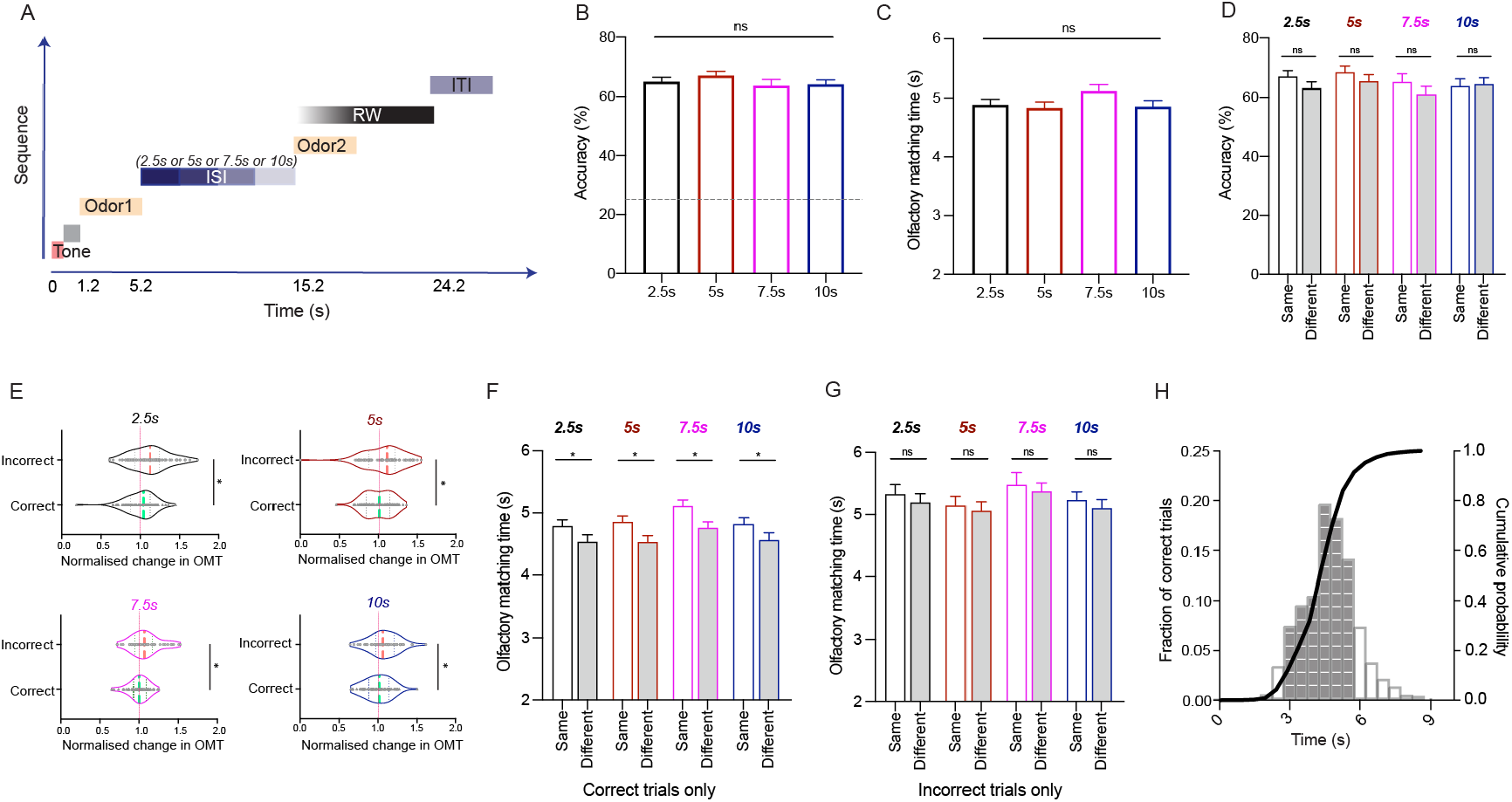
Faster olfactory matching time for correct responses. A. Illustration depicting the sequence of events in a single trial of Olfactory Matching (OM) task over time. Each trial is initiated with a tone for 200ms followed by delivery of odor 1 after 1s of tone presentation. The entire trial may take 11.7s-24.2 s depending on the inter-stimulus interval (ISI) after odor 1 and the olfactory matching time (OMT) after the odor 2 is delivered. (RW refers to Response Window during which the subject decided if the two odors were same or different and thus, initiates a motor response to press on the response console using which OMT is calculated, ITI refers to the Inter-trial interval, is of 5s which marks the end of a trial). B. OM accuracy averaged across subjects (Each subject carried out 20 trials) for varying ISI indicated that humans are able to perform the task at similar accuracy levels across 2.5s, 5s, 7.5s and 10s ISI durations. (N_2.5s_ = 75 sessions (black bar), N_5s_ = 78 (red bar), N_7.5s_ = 53 (green bar) and N_10s_ = 63 (blue bar); p = 0.42, F = 0.93, One-Way ANOVA). Each session consisted of a block of 20 trials of OM task done by a human subject. Dotted line represents the chance level performance set at 25.75%. C. OMT (in seconds) also remains comparable across the OM tasks of varying ISI suggesting that the overall response time remained unperturbed by the time lag between the stimuli, at least, with the durations that we evaluated (N_2.5s_ = 75, N_5s_ = 78, N_7.5s_ = 53 and N_10s_ = 63; p = 0.21, F = 1.524, Ordinary One-Way ANOVA). D. Accuracies for ‘same’ vs. ‘different’ trials, averaged across subjects, showed similar performance levels between the two kinds of trials in the OM task across the four different ISIs (N_2.5s_ = 75, p = 0.17; N_5s_ = 78, p = 0.36; N_7.5s_ = 53, p = 0.24; and N_10s_ = 63, p = 0.85; Paired t-test for comparing same vs different per ISI). E. Normalized changes in OMT are depicted using violin plots. Shift in OMT between correct vs. incorrect trials were observed, suggesting a significantly increased OMT for incorrect responses on the OM task across the four different ISI values. The green dotted line represents the OMT mean of correct while the red dotted line (shifted towards left) represents the mean OMT of incorrect response. The gray dotted line is centered at value 1, which is obtained from calculating the average OMT for the total correct responses for each ISI. OMTs for the corresponding correct and incorrect trials of a particular ISI, were then normalized to this value (p_2.5s_ < 0.0001, p_5s_ = 0.003, p_7.5s_ < 0.0001 and p_10s_ < 0.0001, Paired t-test for each ISI plot). F. Change in OMT between same vs. different correctly responded trials exhibit shorter time taken to decide whether the matched odors were different as compared to the decision pertaining to same correct decisions (p_2.5s_ = 0.0003, p_5s_ < 0.0001, p_7.5s_ = 0.0003 and p_10s_ = 0.0013, Paired t-test for each ISI plot). G. Averaged OMTs are statistically similar for same vs. different trials when the responses were incorrectly chosen by the subjects suggesting OMT as an important read-out for determining certainty and its ability to be modulated while distinguishing same vs. different trials (p_2.5s_ = 0.34, p_5s_ = 0.57, p_7.5s_ = 0.58, and p_10s_ = 0.36, Paired t-test for each ISI plot). H. Histogram representing fraction of correct trials falling across the OMT divided into bins of 500ms each (left y-axis). Shaded region represents ∼80% of the correct responses for which OMT is lying between 3s to 5.5s. A cumulative probability curve (black) is super-imposed on the histogram (right y-axis).

Here, we do observe that the behavioral manifestation of WM is subjected to uncertainty. Accuracy and reaction times in perceptual decision-making tasks are generally derived from the ‘drift-diffusion’ models [25,26]. In this model, human brain is supposed to collect evidences from the stimulus which are perturbed by noise and is accumulated over time until it is enough, at which point, the decision is made [1]. As the neural activity that supports WM is noisy and resource limited, we see that longer responses are often inaccurate. Our results are indeed consistent with the existing understanding of human perception using other sensory modalities [27–29].

In the OM task so far, the accuracy and OMT were tested from trials of equal sensory complexity (i.e., monomolecular odorants). We next investigated WM in binary odor mixture matching task. We selected two different odor mixtures with varying ratios (80-20 vs. 20-80 and 60-40 vs. 40-60) and ensured sharp odor-mixture onset and offset using our PID measurements (Figure 3A). Detection indices were also comparable across 10 binary mixture of odors measured in 107 subjects (Figure 3B) and remained statistically similar to simple monomolecular detection accuracy (Figure 3C; p = 0.1, Unpaired t-test, two-tailed).

**Figure 3:**
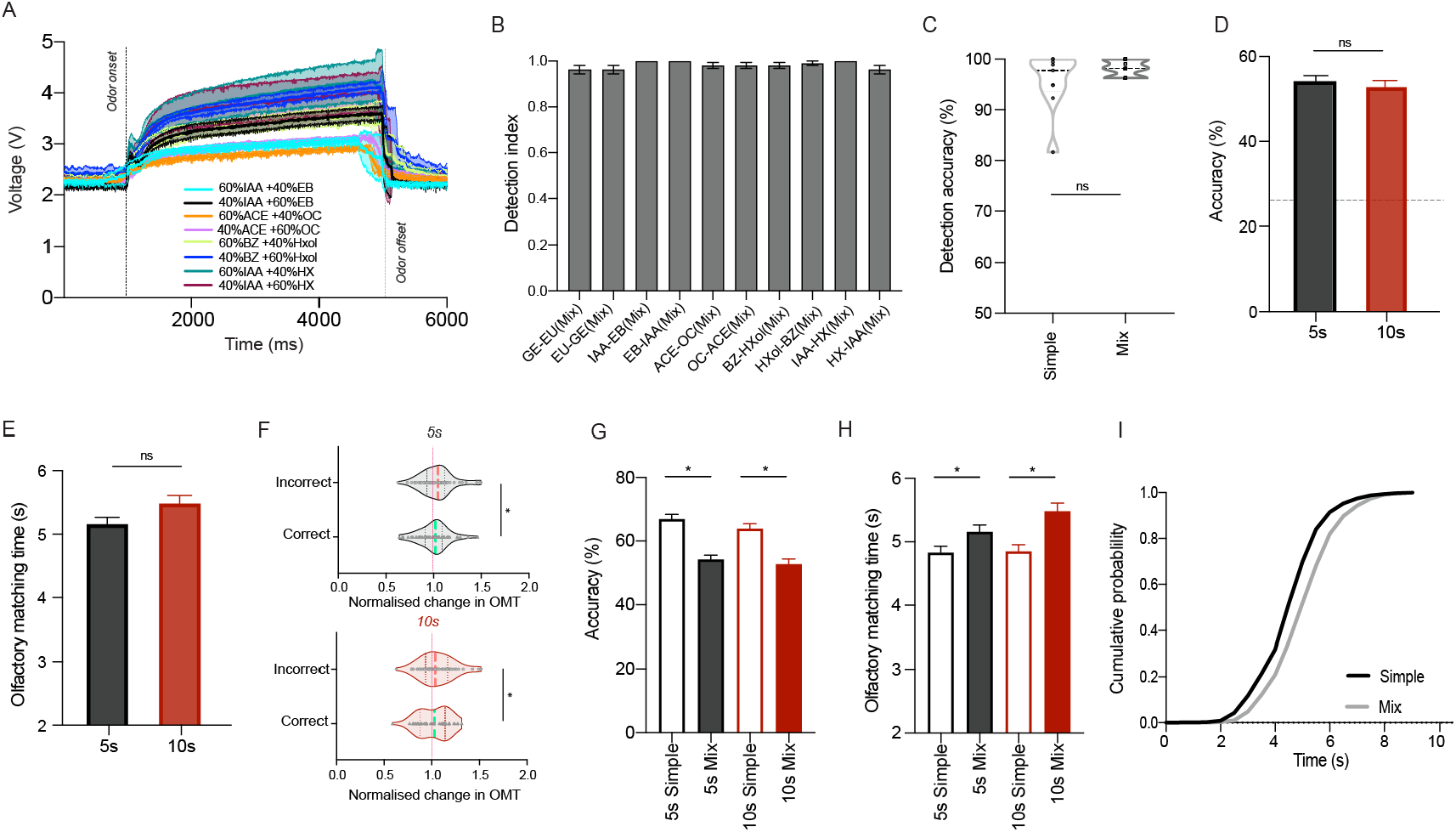
Olfactory matching time is longer for binary-mixture odors. A. PID measurements of changes in voltage amplitudes for binary mixtures (60%-40% or 80%-20%) of odors are plotted (IAA: Isoamyl Acetate, EB: Ethyl Butyrate, ACE: Acetophenone, OC: Octanal, BZ: Benzaldehyde, Hxol: Hexanol, HX: Hexanal). B. Detection indices of binary-mixtures of odors delivered via the OAM, averaged across subjects (N = 107 subjects). C. The detection accuracy calculated from their DIs reveal similar performance of detecting monomolecular (simple) and binary mixtures of odors (N_simple odors_ = 9 odors, N_mix odors_ = 10 odors, p = 0.1, Unpaired t-test). D. OM accuracy averaged across subjects for 5s and 10s ISI indicated that humans are able to perform the task at similar accuracy levels for binary-mixture tasks for the two ISIs (N_5s_ = 58 subjects, N_10s_ = 47; p = 0.48, Unpaired t-test, two-tailed). Chance level performance for a binary mixture OM task was calculated to be at 25.04% E. Overall OMT (including both correct and incorrect trials) averaged across subjects are comparable for 5s and 10s ISI groups (N_5s_ = 58, N_10s_ = 47; p > 0.05, Unpaired t-test, two-tailed). F. Normalized change in OMT is shifted towards right (i.e. it is longer) for the incorrect trials in case of both 5s and 10s ISI groups for binary-mixture OM tasks as compared to the correct trials. The black dotted line is centered at value 1, which is obtained from calculating the average OMT for the total correct responses for each ISI. OMTs for the corresponding correct and incorrect trials of a particular ISI, were then normalized to this value (N_5s_ = 58, p_5s_ = 0.0092 and N_10s_ = 47, p_10s_ = 0.013, Paired t-test). G. OM performance levels as quantified using the averaged OM accuracy for 20 trials per subject revealed better performance for OM task of monomolecular odors (simple) as compared to binary-mixtures, for both 5s and 10s (p < 0.0001 for 5s-simple vs. 5s-mixture as well as for 10s-simple vs. 10s-mixture, Unpaired t-test, two-tailed). H. OMT was longer for binary mixture tasks, which correlates with decreased accuracy, for both, 5s and 10s ISI (p = 0.028 for 5s-simple vs. 5s-mixture and p = 0.0002 for 10s-simple vs. 10s-mixture, Unpaired t-test, two-tailed). I. Cumulative probability distribution of correct responses over the OMT for simple vs. mixture depicts significantly longer OMT is needed for accurately performing the mixture OM task (p < 0.0001, Kolmogorov-Smirnov test).

Although no overall observable differences existed in performance accuracies across different ISIs in the monomolecular OM task, individual feedback from the participants disclosed the difficulty to remember and recollect when ISI was set at 10s. Thus, for odor mixture tasks, we chose 10s ISI to test if human perception is shifted in the either direction, in such a difficult task. Along with 10s ISI, 5s ISI was also chosen wherein most of the subjects found it easy to perform the monomolecular OM task. The OM accuracy, as was observed in the simple task, remained comparable between the two ISI used (5s and 10s) even in a complex mixture matching task (Figure 3D; N_5s_ = 58 subjects, N_10s_ = 47; p = 0.48, Unpaired t-test, two-tailed; Supplementary Figure 2A; p > 0.1, Paired t-test for same vs. different trials across the two ISIs). The OMT was also comparable between 5s and 10s ISI in this task (Figure 3E; p > 0.05, Unpaired t-test, two-tailed) suggesting that the limit of WM, both, in terms of reliably holding the information to a certain time limit and the complexity of odor stimuli could be beyond the ISIs and complexity that we utilized in our study. As in the simple odor OM task, the OMT showed a rightward shift for incorrect trials for the two ISIs that we tested in the binary mixture task (Figure 3F; p_5s_ = 0.0092 and p_10s_ = 0.013, Paired t-test).

Interestingly, humans performed the simple task more accurately and with a faster OMT as compared to the mixture task suggestive of an increased uncertainty in their responses when challenged with a complex task (Figure 3G and 3H; p < 0.05 across all measurements, Unpaired t-test, two-tailed). Accuracy remained comparable across two ISIs for same vs. different binary mixture trials (Supplementary Figure 2A). As was observed for simple matching task, the OMT was significantly longer for correctly matched binary mixture similar trials at 5s ISI (Supplementary Figure 2B; p_5s_ = 0.013, Paired t-test). However, it remained statistically similar at the longest delay of 10s ISI (Supplementary Figure 2B; p_10s_ > 0.1, Paired t-test). This is indicative of the increased complexity posited by 10s ISI and the complexity of odors effecting concurrently in a 10s ISI mixture task As expected, the incorrectly matched binary mixture trials remained statistically similar whether the trials were same or different (Supplementary Figure 2C, p > 0.1, Paired t-test). Upon comparing the correctly matched simple vs. mixture trials across different ISI conditions and the trial types (same or different), we consistently found higher OMT for binary odor mixture trials (Supplementary Figure 2D; p < 0.05 for all comparisons, Unpaired t-test). This is also shown in a cumulative plot, which proved that binary mixture tasks require significantly longer OMT to correctly respond in an OM task as compared to simple tasks (Figure 3I; p < 0.0001, Kolmogorov-Smirnov test and Supplementary Figure 2E). Indeed, OMT for mixture task was lying between 4s to 6.5s for approximately 77% of the correct responses (Supplementary Figure 2E).

Our results show that although the WM performance in an OM task is unaffected across different ISI conditions we tested, it is challenged upon increasing the stimulus complexity. Also erroneous responses, regardless of the time delay or the stimulus complexity, were undertaken by the humans when taking longer OMT to respond. This suggests that uncertainty increases when OMT is longer. Finally, upon comparing the females vs. males for the two parameters, accuracy and OMT, we observe comparable WM capabilities suggesting no effect of gender on the olfactory fitness in humans (Supplementary Figure 3A and B; N_males_simple_ = 144 subjects, N_females_simple_ = 125, N_males_mix_ = 49, N_females_mix_ = 56; p > 0.05, Unpaired t-test, two-tailed).

Olfactory perceptual space has been characterized using multiple human psychophysical experimental approaches [30–32]. Working memory based tasks are also used to study perceptual similarities in rodents [33]. Indeed, various approaches resulted in contrasting findings on how olfactory stimulus similarity affects response times [34–40]. Our results clearly demonstrate that matching of similar binary odor mixture takes longer compared to monomolecular odors. Hence, allowing subjects to match complex odor stimuli and taking decisions without any time constraints, proved that accurate complex decisions happen at a temporal cost.

Bi-directional connectivity is proposed between the primary olfactory cortical areas and the ones encompassing the olfactory-thalamic cortical route [41]. The Mediodorsal nucleus of the Thalamus (MDT) receives inputs from several olfactory structures including Piriform Cortex (PCx), Olfactory Tubercle (OT), Cortical Amygdala and the Lateral Entorhinal Cortex. MDT is known to modulate attention in olfactory task and projects to Orbitofrontal cortex (OFC). fMRI based studies have displayed enhanced functional connectivity between PCx-MDT and MDT-OFC during an olfactory attention task [42]. In rodents, olfactory tubercle receives direct input from medial prefrontal cortex, which plays a central role in selective attention to odors [43]. Attention may play a significant role in completing olfactory matching task with high accuracy especially when the subjects are challenged with longer ISIs. However, we do not know the maximum duration of olfactory information in working memory, which is normally 10-15s without any repetition, a reported duration from other sensory systems [44].

Entorhinal cortex (EC), a region in the medial temporal lobe primarily receives projections from hippocampus, playing a role in memory. Within olfactory areas, it has bi-directional connections from OB and PCx [45]. This region has been shown to maintain cue-specific neuronal activity in a delayed working memory task [46]. The human parietal cortex, that gathers information from EC may also facilitate orienting the attention of the subject during a decision-making task [47]. Focusing on the OB-PCx-EC pathway may help in delineating the circuit mechanisms of ‘certain’ decisions. However, more experiments are needed to find out maximum duration of olfactory working memory under varying stimulus complexity and thereby to study the limits of olfactory deliberation. In summary, our findings will facilitate probing WM-dependent changes in the olfactory perception and thereby, cognitive fitness in individuals with neurodegenerative and neurological disorders under infectious and non-infectious conditions.

## METHODS

### Key resources table

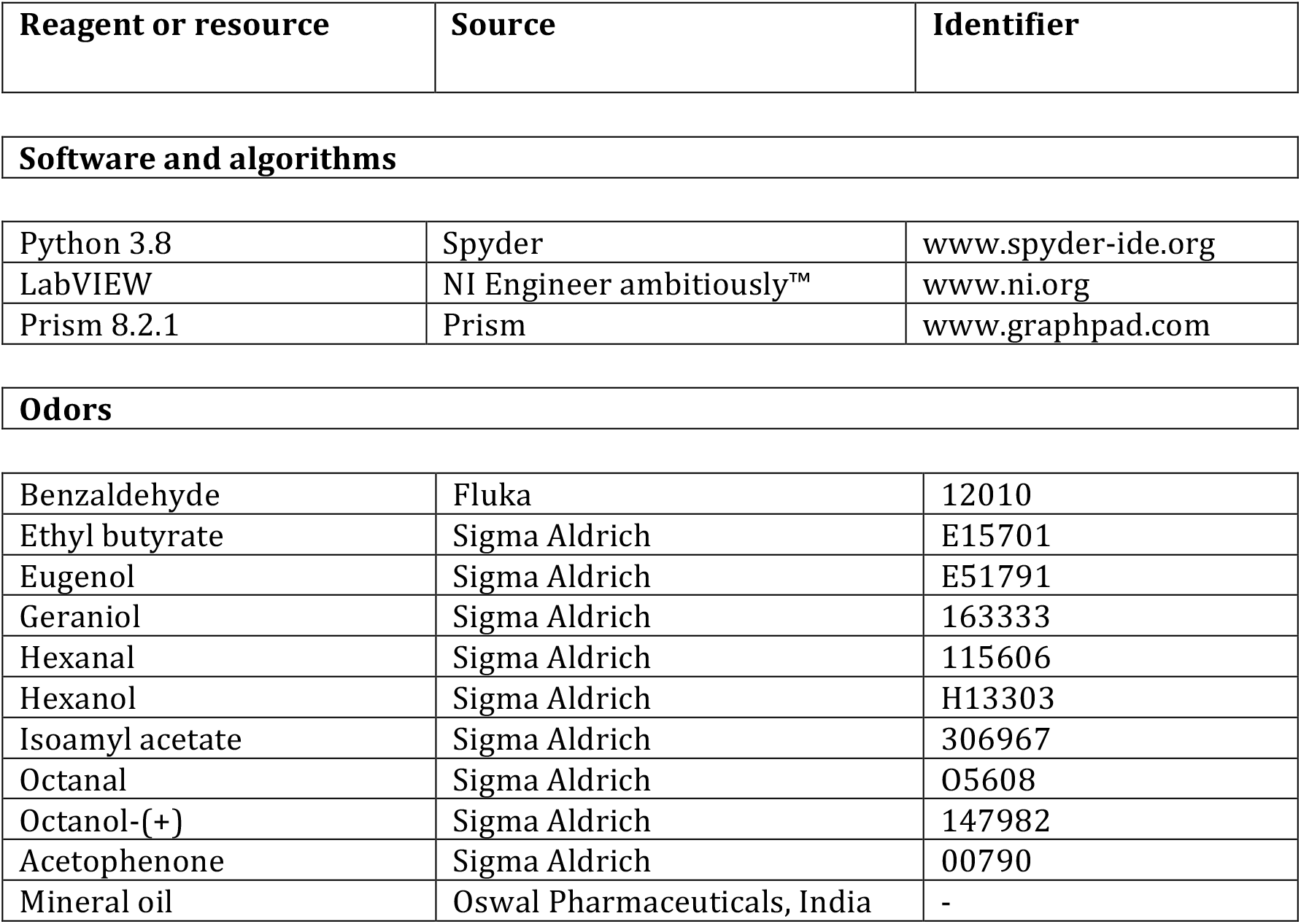

| Reagent or resource | Source | Identifier |
| --- | --- | --- |
| Software and algorithms |  |  |
| Python 3.8 | Spyder | www.spyder-ide.org |
| LabVIEW | NI Engineer ambitiously™ | www.ni.org |
| Prism 8.2.1 | Prism | www.graphpad.com |
| Odors |  |  |
| Benzaldehyde | Fluka | 12010 |
| Ethyl butyrate | Sigma Aldrich | E15701 |
| Eugenol | Sigma Aldrich | E51791 |
| Geraniol | Sigma Aldrich | 163333 |
| Hexanal | Sigma Aldrich | 115606 |
| Hexanol | Sigma Aldrich | H13303 |
| Isoamyl acetate | Sigma Aldrich | 306967 |
| Octanal | Sigma Aldrich | O5608 |
| Octanol-(+) | Sigma Aldrich | 147982 |
| Acetophenone | Sigma Aldrich | 00790 |
| Mineral oil | Oswal Pharmaceuticals, India | - |

### Resources availability

Further information and requests for resources should be directed to, and will be fulfilled by the Lead Contact, Nixon M. Abraham. Materials availability [17]

### Subject details

A total of 394 human subjects (197 males and 197 females) belonging to the Indian subcontinent participated in this study. 143 subjects carried out the questionnaire-based assessment of olfactory detection indices for the commonly available odorous objects. To closely evaluate the features of olfactory matching skills of humans, Olfactory-action meter or OAM [16,17] was employed. 256 (132 males and 124 females) of the total subjects participated in the OAM based olfactory detection and Olfactory matching (OM) evaluation across different ISIs and stimulus complexity (i.e simple and binary-mixture odors). Either one or two sessions were carried out. Participants signed a consent form before taking part in the experiment.

## Method details

### Online questionnaire for odorous objects detection

This test was carried out by participants in their households using commonly available odorous objects. Data was collected using an online questionnaire to probe the general smelling abilities of humans. For each subject, Detection index value of ‘1’ indicated successful detection while ‘0’ meant failed at detecting the object using olfaction. Odorous items were divided into Clusters; Cluster 1 composed of ‘Citrus fruits’ (Yellow bars), Cluster 2 had ‘Odorous food items’ (Brown and Green bars), Cluster 3 encompassing flowers, talcum and deodorants/soaps (Light green bars), Cluster 4 consisted of ‘fruits other than citrus’ (Orange bars), Cluster 5 included ‘spices’ (Maroon bars) and Cluster 6 consisted of ‘Fats and Oils’ (Olive colored bars) (See Figure 1A). Data from participants who reported prior loss or reduction in sense of smell was not taken into consideration.

### Olfactory action-meter (OAM)

The olfactory-action meter is a ten-channel olfactometer that delivers odors with high temporal precision, controlled by an automated paradigm (Figure 1B) [17]. The apparatus delivers odorized air through a 12-channel glass funnel connected to a ventilator mask, where subjects inhaled various odors. The deodorized air was pumped into the olfactometer at a rate of 5 liters per minute using an air pump. Using a metallic custom-built manifold, the air inside the olfactometer was split into eleven channels. These eleven channels were coupled with ten mini and one large mass flow controller (MFC) (Pnucleus Inc.). Functioning of MFCs were controlled by custom written LabVIEW scripts, which allowed the experimenter to control the onset of airflow and its rate from each MFC. Upon initiation of the trial, the odor bottle received air input from mini MFCs. Simultaneously, the output from the main (MFC) was routed to 10 channels via an array of solenoid valves, with one solenoid valve corresponding to one odor channel. The output air from the valve-controlled channel merged with the odorized air from the odor bottle before entering the glass nozzle connected to a ventilator mask that subjects wore during the course of experiment.

### Odor preparation and delivery

The stimulus profiling for monomolecular odors (2% dilution in mineral oil) as well as binary-mixture odors (1% dilution in mineral oil at 60%-40% or 80%-20% concentrations of the odors) were measured using miniPID acquired from Aurora Scientific (Figure 1C and 3A). All odors were purchased from Sigma-Aldrich/Fluka and had purity >90% (details in Key resources table). All the PID measurements were done while odor stimulus was provided for 4s, which was the stimulus duration used during odor detection and matching tasks.

### Odor detection task

During the odor detection task, odors were sequentially delivered to the individuals using the OAM (Supplementary Figure 1). They were asked to report (a Yes or a No) whether they detected the odor. Successful odor detection response was recorded as 1, whereas the wrong response was recorded as 0. Each odor was delivered for 4s, and the sequence of the odors was randomized across subjects. During a session, each subject conducted a detection task for either simple odors or binary-mixture odors. The detection scores for each odor were averaged across all individuals and referred to as the detection index for that odor. Overall detection abilities were evaluated using detection accuracy, which was determined by the percentage of odors detected.

### Olfactory matching (OM) task

Participants that performed olfactory detection task were further requested to participate in the olfactory matching task as well (Supplementary Figure 1). Each session of the OM task constituted 20 trials. The trial began with a tone of 200ms, followed by the first odor after 1s. The first odor was delivered for 4s, followed by an inter-stimulus (ISI) delay, and then the second odor was delivered for 4s. In a trial, the two odors provided could be the same or different, and participants were asked to compare two odors and decide whether the odors were ‘same or ‘different’ by pressing respective keys on a response console. Response console also had a ‘Repeat’ button that allowed participants to repeat the trial in case of ambiguity in the decision. The subjects had to respond by pressing the button within the response window (9s) that began with the onset of the second odor (4s) and lasted 5s after its offset. The subsequent trial started after an inter-trial interval of 4s of making the decision (i.e., pressing response buttons on the console). Olfactory matching abilities were measured for simple odors as well as binary-mixture odors in our study. Based upon the odors detected by the subjects, odor pairs for the OM task were generated. Maximum of 5 odor pairs were provided during a session. Olfactory matching task was done while participants encountered differing ISI (2.5s, 5s, 7.5s and 10s for simple odors and 5s and 10s for binary-mixture odors), wherein ISI among trials during a particular session was kept same (Figure 2A). Olfactory matching performance was analyzed by calculating the percentage of correct trials in a session. Chance level performance in the OM task was calculated using binomial expansion:

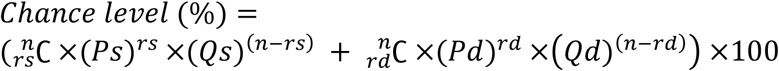

where *n* = total trials, *rs* = number of same trials, *rd* = number of different trials, *Ps* = Probability of carrying the same trial correctly, *Pd* = Probability of carrying the different trial correctly, *Qs* = Probability of carrying the same trial incorrectly and *Qd* = Probability of carrying the different trial incorrectly.

The ‘Same’ and ‘Different’ probabilities are added which reveals the probability of carrying out correct trial at a chance, thus, referring to it as the chance level accuracy. This was found out to be ∼25% for the OM task.

## Data and statistical analysis

Data analysis was performed by custom written python scripts. All statistical analyses were done using GraphPad Prism 8. We used Two-tailed paired/unpaired t-tests, Kolmogorov-Smirnov test, Analysis of variance (ANOVA) and associated post-hoc tests.

## Author contributions

N.A.: Conceptualized the study and designed the experiments.

R.B., M.P., S.M., A.B., S.K., S.P., T.M., E.M.G., P.S., and S.D.M: Collected the data

S.M., M.P. and R.B.: Analyzed the data.

M.P., S.M. and N.A.: Wrote the manuscript with comments from other authors.

## Ethics committee approval information

All experimental procedures and protocols used in this study were approved by the IISER Ethics Committee for Human Research.

## Acknowledgements

We thank Laboratory of Neural Circuits and Behaviour (LNCB) members for fruitful discussions and comments on the Manuscript. We thank the IISER community for the support extended during the data collection.

This work was supported by the DST-Cognitive Science Research Initiative (DST/CSRI/2017/271 to N.A.), DBT/Wellcome Trust India Alliance intermediate grant (IA/I/14/1/501306 to N.A.), CSIR Fellowships (R.B., S.M., and S.D.M.), UGC NET Fellowship (A.B.) and IISER-Pune Fellowship (M.P.).

