## Supplementary Figures for "Uncertainty revealed by delayed responses during olfactory matching"

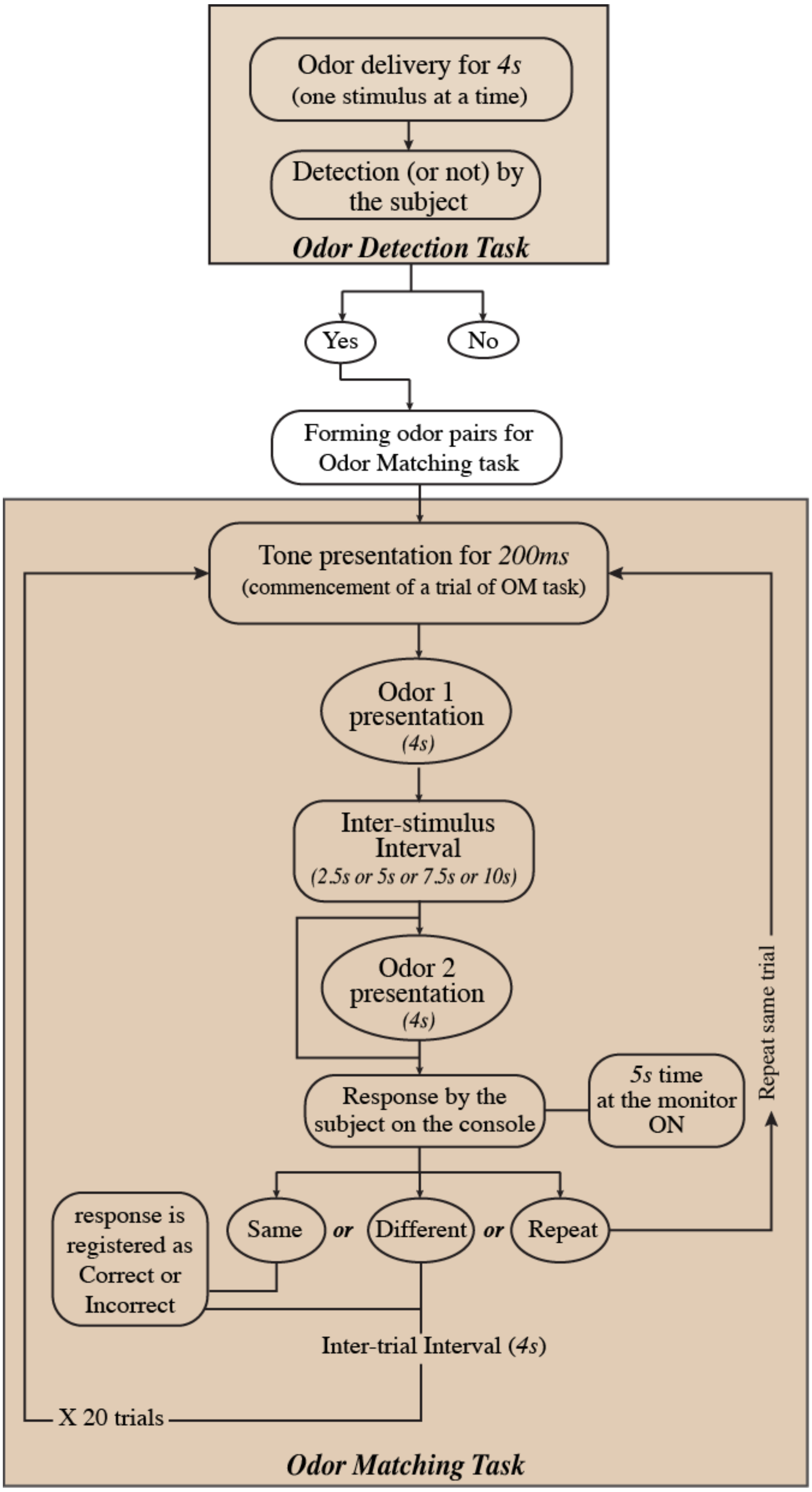


**Figure S1: Flowchart of the Odor Detection and Odor Matching task**

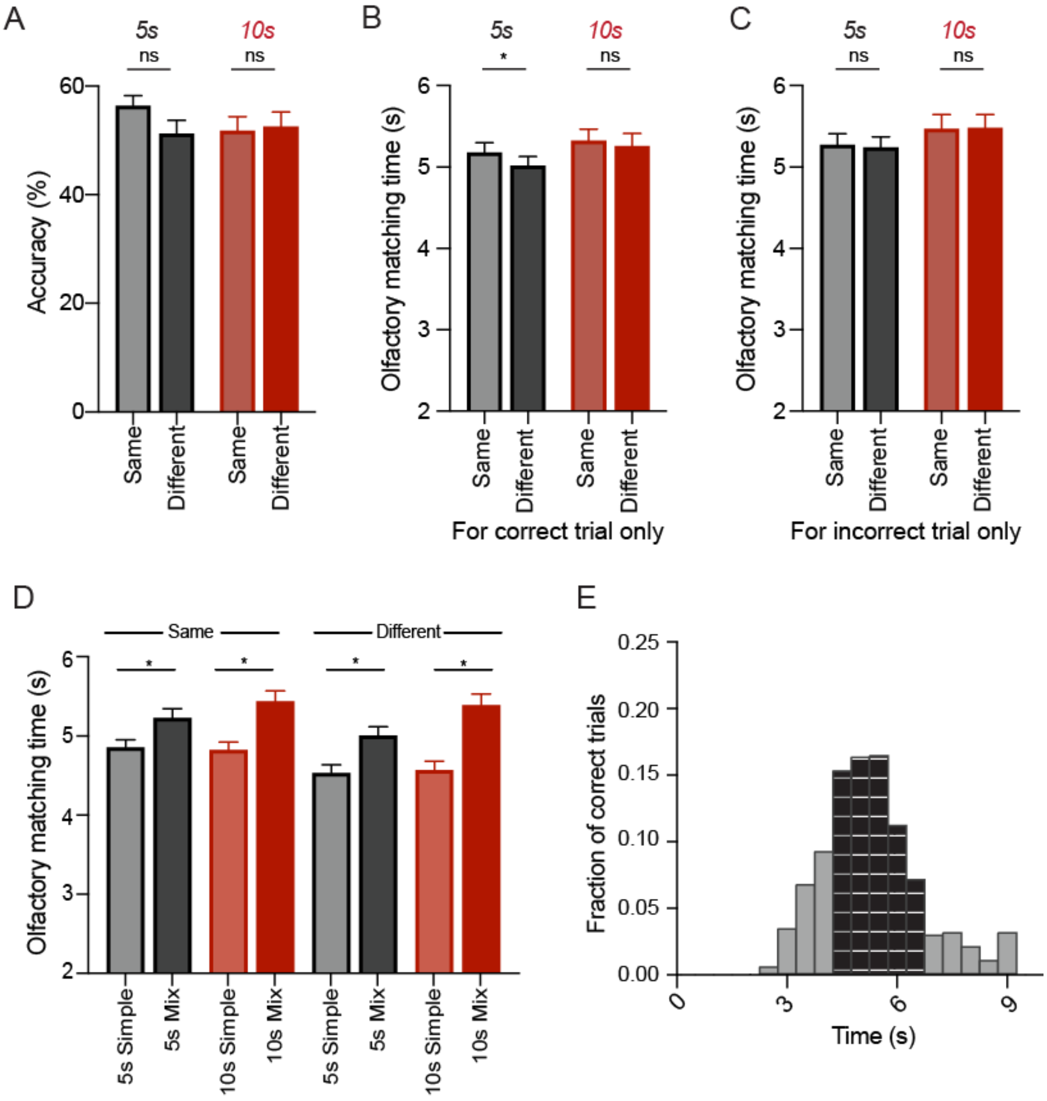


**Figure S2: Similar performance levels at longest time-delay, binary mixture OM task between the two kinds of responses**

A. Accuracies for 'same' vs. 'different' trials, averaged across subjects, shows similar performance levels between the two kinds of trials in the binary mixture OM task across the two different ISIs ($N_{5s}$ = 58, $p_{5s}$ = 0.13 and $N_{10s}$ = 47, $p_{10s}$ = 0.86, Paired t-test).

B. Change in OMT between same vs. different correctly responded trials exhibit shorter time taken to decide whether the matched odors were different as compared to the decision pertaining to same correct decisions only for 5s ISI while they are comparable for 10s ISI duration ($p_{5s}$ = 0.013 and $p_{10s}$ = 0.6, Paired t-test for each ISI plot).

C. Averaged OMTs are statistically similar for same vs. different trials when the responses are incorrectly responded by the subjects for both the ISI durations $p_{5s}$ = 0.81 and $p_{10s}$ = 0.91, Paired t-test for each ISI plot).

D. Comparison of same vs. different trials OMT (in seconds) between simple and binary mixture tasks indicates that humans take longer OMT to correctly match mixture trials across both 5s and 10s ISIs ($p_{5s} = 0.0161$ and $p_{5s} = 0.0025$ for same and different correct trials between simple vs. mixture tasks respectively when ISI is 5s, $p_{10s} = 0.0003$ and $p_{10s} < 0.0001$ for same and different correct trials between simple vs. mixture tasks respectively when ISI is 10s, Unpaired t-test for all comparisons).

E. Histogram representing fraction of correct trials falling across the OMT divided into bins of 500ms each (left y-axis) for binary mixture OM task. Shaded region represents ~77% of the correct responses for which OMT is lying between 4s to 6.5s.

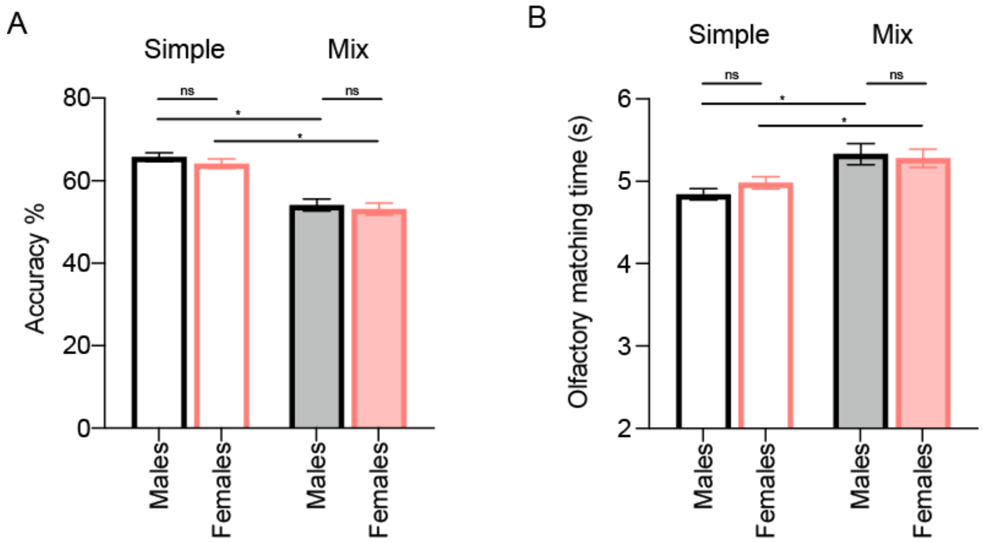


**Figure S3: Similar performance levels between males and females in a simple and binary mixture OM task**

A. Accuracies are comparable between males and females for simple and mixture OM task ($N_{males\_simple} = 144$ subjects, $N_{females\_simple} = 125$, $N_{males\_mix} = 49$, $N_{females\_mix} = 56$; $p = 0.32$ for simple and $p = 0.64$ for mixture, Unpaired t-test, two-tailed). Between the simple and mixture tasks, performance levels decrease as odor complexity increases in both male and female human subjects ($p < 0.0001$, Unpaired t-test, two-tailed for both).

B. Average OMT are comparable between males and females for simple and mixture OM task ($N_{males\_simple} = 144$ subjects, $N_{females\_simple} = 125$, $N_{males\_mix} = 49$, $N_{females\_mix} = 56$; $p = 0.16$ for simple and $p = 0.76$ for mixture, Unpaired t-test, two-tailed). Between the simple and mixture tasks, OMT increases as odor complexity increases in both male and female human subjects ($p = 0.0006$ for males and $p = 0.026$ for females, Unpaired t-test, two-tailed for both).
